# The collapse of the spindle following ablation in S. pombe is mediated by microtubules and the motor protein dynein

**DOI:** 10.1101/2020.10.20.347922

**Authors:** Parsa Zareiesfandabadi, Mary Williard Elting

## Abstract

A microtubule-based machine called the mitotic spindle segregates chromosomes when eukaryotic cells divide. In the fission yeast *S. pombe*, which undergoes closed mitosis, the spindle forms a single bundle of microtubules inside the nucleus. During elongation, the spindle extends via antiparallel microtubule sliding by molecular motors. These extensile forces from the spindle are thought to resist compressive forces from the nucleus. We probe the mechanism and maintenance of this force balance via laser ablation of spindles at various stages of mitosis. We find that spindle pole bodies collapse toward each other following ablation, but spindle geometry is often rescued, allowing spindles to resume elongation. While this basic behavior has been previously observed, many questions remain about this phenomenon’s dynamics, mechanics, and molecular requirements. In this work, we find that previously hypothesized viscoelastic relaxation of the nucleus cannot fully explain spindle shortening in response to laser ablation. Instead, spindle collapse requires microtubule dynamics and is powered at least partly by the minus-end directed motor protein dynein. These results suggest a role for dynein in redundantly supporting force balance and bipolarity in the *S. pombe* spindle.

**STATEMENT OF SIGNIFICANCE:** *S. pombe* serves as an important model organism for understanding cell division. Its structurally simple mitotic spindle is especially suited for mechanical perturbation. Since *S. pombe* undergoes a process of closed cell division, without breakdown of the nuclear envelope, force may be exerted between its nuclear envelope and spindle. Here, we mechanically sever spindles via laser ablation to probe this force balance. Following ablation, *S. pombe* spindle fragments collapse toward each other. We find that, contrary to prior expectations, forces from the chromosomes and nuclear envelope are not responsible for this collapse. Instead, it is microtubule-dependent, and is powered at least in part by the minus-end directed microtubule motor protein dynein.

## INTRODUCTION

The mitotic spindle, a microtubule-based cellular machine, is responsible for accurate chromosome segregation in eukaryotes. During mitosis, spindle microtubules attach to the chromosomes, align them, segregate them, and physically deliver them to the two new daughter cells, ensuring each one has exactly one copy of the genetic information of the cell encoded in the chromosomes. Accurate chromosomes segregation is an essential function for life. Mis-segregation leads to aneuploidy, a condition of extra or missing chromosomes, associated with developmental defects and cancer in multicellular organisms [1, 2]. The mitotic spindle machinery is highly conserved across eukaryotes, likely due to its critical function.

*S. pombe* is a eukaryotic model system that is well-established for probing the microtubule cytoskeleton, including the mitotic spindle [3, 4]. Its geometrically simple spindle structure, which comprises a single bundle of microtubules [5], also makes it ideal for mechanical perturbation. *S. pombe* undergoes closed mitosis, with its nuclear envelope remaining intact during chromosome segregation [6, 7]. The timing of mitosis is highly uniform from cell-to-cell and can be divided into three distinct phases defined by the spindle length and rate of elongation [8]: phase one, which includes prophase and spindle formation; phase two, which includes metaphase and anaphase A, during which spindle length remains constant at ~ 2.5 *μ*m; and phase three, which includes anaphase B, during which the spindle extends at a constant rate, elongating the nucleus into a dumbbell shape before it ultimately divides into two daughter nuclei.

Mitotic spindles are built of microtubule protein filaments oriented in the spindle to form a bipolar structure. During *S. pombe* mitosis, microtubule minus ends are anchored at specialized microtubule-organizing centers (MTOCs) called spindle pole bodies (SPBs), while microtubule plus ends extend into both the cytoplasm and nucleoplasm [7, 9]. The cytoplasmic population forms the astral microtubules, which help orient the spindle inside the cell [10–12]. On the other hand, the spindle microtubules remain entirely in the nucleus, anchored at the two SPBs that are embedded in the nuclear envelope throughout mitosis [13]. Three classes of spindle microtubules have been identified in *S. pombe*: those that interdigitate in the mid-zone between the SPBs; those that emanate from one SPB and terminate at the other; and those that end at the kinetochore [5]. Together, these microtubules and their associated proteins form a bipolar structure that ensures robust and symmetric chromosome segregation.

Past work has hypothesized that spindle elongation in *S. pombe* is driven by sliding apart the interdigitating antiparallel microtubules at the spindle’s mid-zone, with involvement by several motor proteins [5, 14–19]. Previous work directly tested this model by mechanically severing spindles via laser ablation, demonstrating that microtubules within the spindle (rather than astral microtubules exterior to the nucleus) generate the force of spindle elongation [11, 12]. How mechanical force-balance is maintained amongst spindle microtubules during *S. pombe* mitosis is not fully clear. Importantly, these previous studies found that, following laser ablation, severed spindle fragments responded by collapsing toward each other. That work suggested that the collapse might be caused by the viscoelastic relaxation of the nuclear envelope and other materials inside the nucleus such as chromosomes, which deform in response to the extensile forces of spindle elongation [11, 12]. However, this model has not yet been tested, nor has the molecular basis of the collapse response been found.

While viscoelastic deformation is one possible explanation for the collapse of *S. pombe* spindles in response to laser ablation, there is also a potential role for active force generation rather than passive relaxation. Indeed, force balance in the spindle in all eukaryotes requires the balancing of extensile forces, usually provided by plus-end directed microtubule motors that slide microtubules apart, and compressive forces, provided by minus-end directed motors or the chromosomes or, in the case of closed mitosis, by the nuclear envelope [20–22]. In many higher eukaryotes, the minus-end directed motor dynein is essential for maintaining spindle force balance and spindle pole integrity [23, 24]. Dynein is not required for mitotic spindle function in *S. pombe* [25], but it does seem to contribute to chromosome biorientation and bundling microtubules at the SPBs [26, 27]. Thus, its mitotic role is not fully understood.

Here, we probe the physical mechanism of the collapse of the *S. pombe* spindle in response to ablation. We show that the viscoelastic deformation of the nucleus does not explain this phenomenon, and instead find a role for microtubule-based force generation. We find that dynein is at least partially responsible for reconnecting the poles of ablated spindles and pulling them toward each other. These results suggest a possible redundant role for dynein in ensuring mechanical force balance in the spindle.

## MATERIALS AND METHODS

### Strains, cell culture, and treatment with small molecule inhibitors

Fission yeast *Schizosaccharomyces pombe* strains used in this study are listed in Table S1. We followed standard growth conditions and media for all fission yeast cultures [28]. We grew all strains on YE5S agar plates at 25 °C before starting liquid cultures. We grew liquid cultures at 25 °C with shaking by a rotating drum. For imaging, we grew MWE16 and MWE23 in YE5S liquid media for 12-24 hours, washed three times using EMM5S, and further grew for 6-18 hours in EMM5S liquid media before imaging. We grew MWE2 and MWE10 in YE5S media for 12 to 24 hours before imaging.

For treatment with methyl 2-benzimidazole carbamate (MBC; Sigma 45368-250M), we washed the cell cultures twice with the same growth media at the appropriate concentration of MBC, prepared from a 20 mM stock solution in DMSO that was stored at −20 ° C. We added latrunculin A (LatA; EMD Millipore) directly to the cell culture from a stock solution of 20 mM in DMSO. For DMSO control data (Fig. 4 and 5), we washed the cells twice with DMSO control media, which we prepared by adding 5ul of DMSO to 2ml of YE, to match the final concentrations of DMSO in the MBC media described above. We conducted laser ablation experiments within 2-4 minutes of drug or DMSO addition.

**FIGURE 1.**
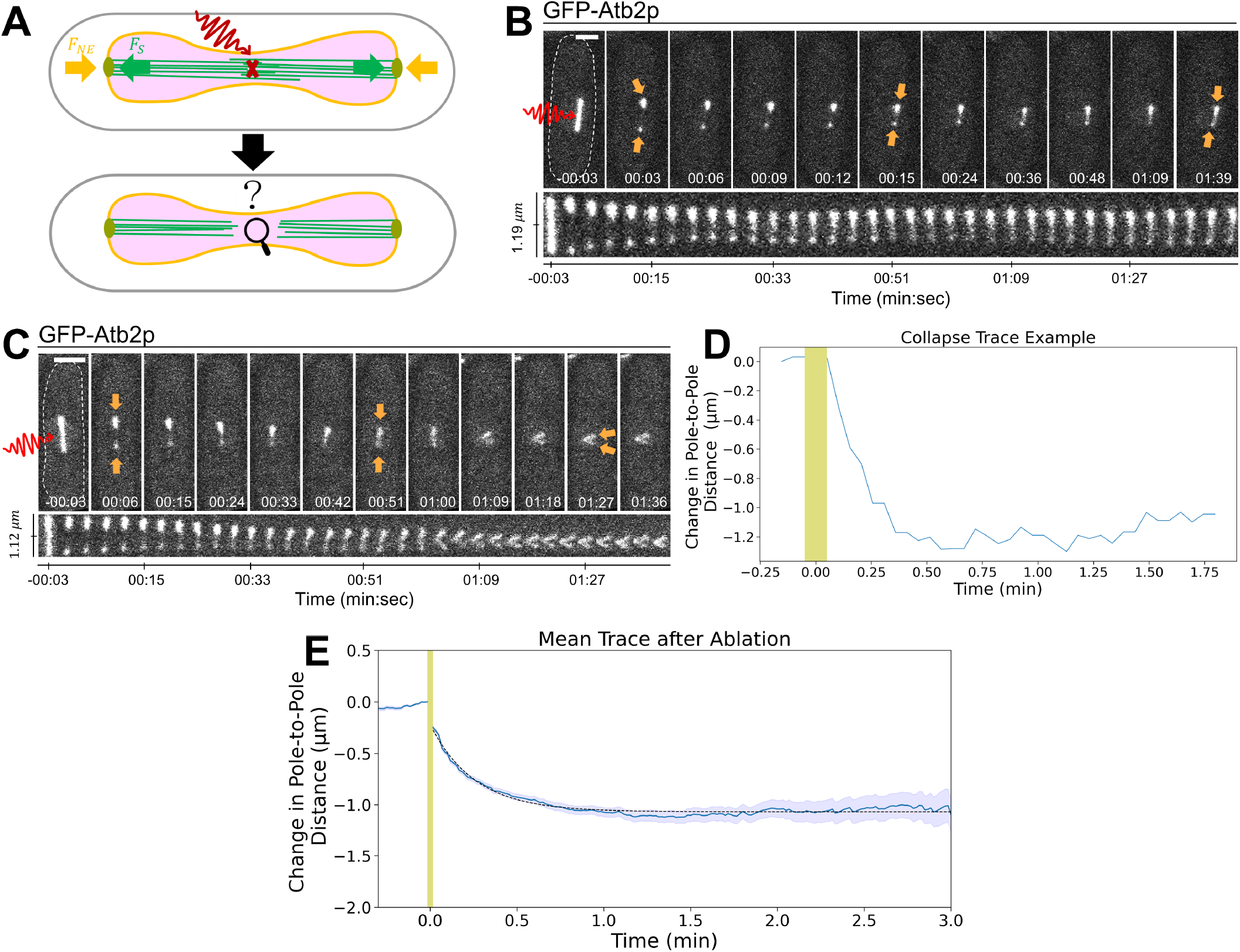
Response of the *S. pombe* mitotic spindle to laser ablation. (A) Experimental schematic. We target the spindle microtubules (green) in the *S. pombe* nucleus (magenta) for laser ablation (red X). We track the response of the spindle, the nuclear envelope (orange), and the spindle pole bodies (lime). Arrows indicate the presumed compressive force of the nuclear envelope, opposed by the extensile force of spindle elongation. (B) and (C) Two examples of *S. pombe* GFP-Atb2p spindles ablated near the mid-region during phase two of spindle elongation. Spindle ends (orange arrows) collapse toward each other after ablation. Top, highlighted frames; below, a montage of all frames. Dashed line in the first frame, cell boundary. Scale bars, 2 *μ*m. Time, min:sec. (D) Change in the pole-to-pole distance over time for the example in (B). (E) The mean trace of change in pole-to-pole distance over time for *n* = 177 ablated spindles (solid blue line) and s.e.m. for all traces (blue shaded region). Black dotted line is a fit to an exponential, with amplitude 0.84 0.02 *μ*m and time constant 0.25 0.01 min. We cannot image during ablation, and the yellow shaded regions in (D) and (E) indicate this period.

**FIGURE 2.**
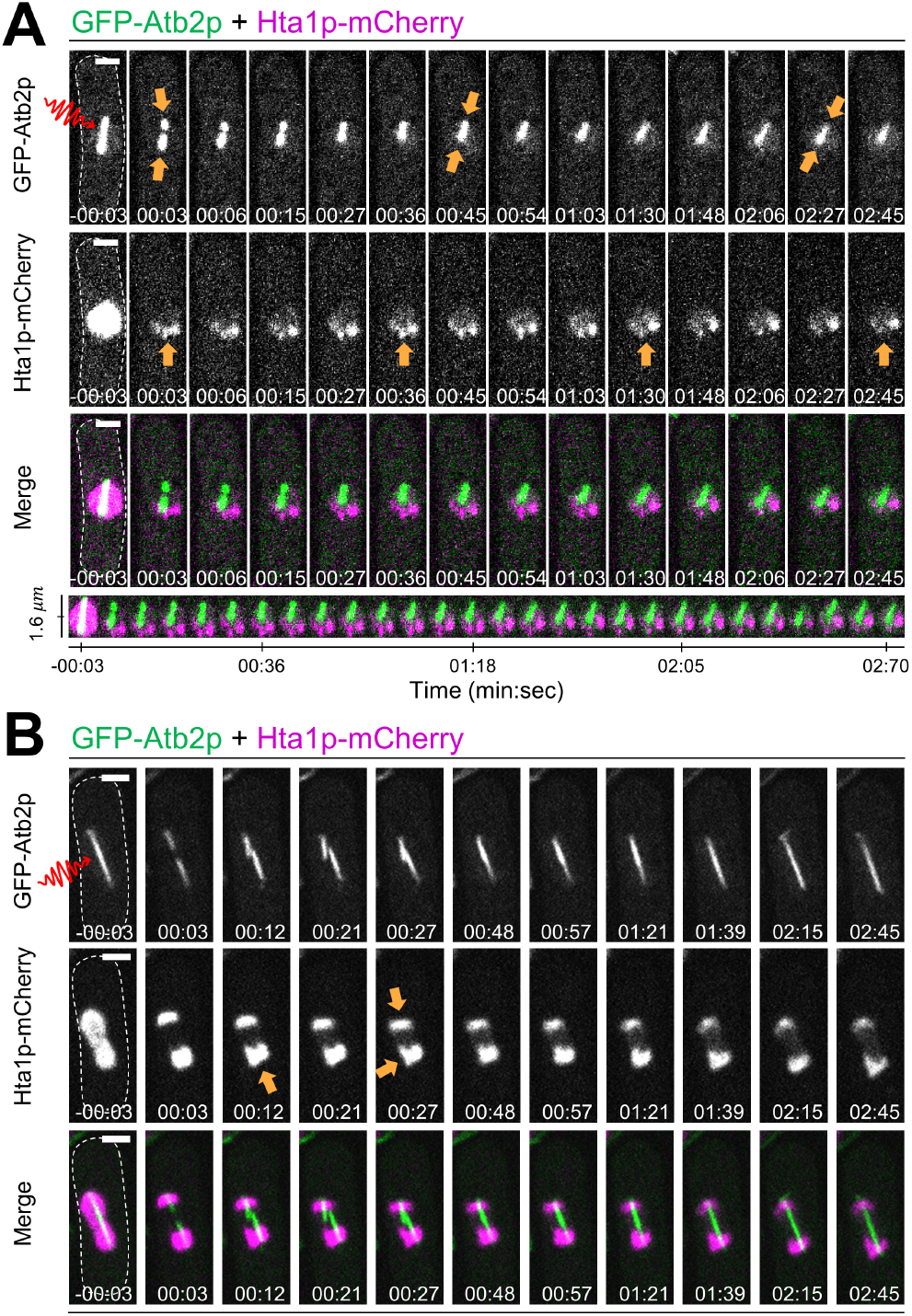
Response of histones to spindle laser ablation. (A) and (B) Two examples of *S. pombe* spindles expressing GFP-Atb2p plus Hta1p-mCherry and ablated near the mid-region during early phase three and mid-phase three of spindle elongation, respectively. Arrows point to spindle ends and histones, indicating the inward collapse of the spindle and histones’ movement after ablation. A montage of collapse is displayed beneath for (A). The spindle collapses inward in both examples. Scale bars, 2 *μ*m. Time, min:sec.

**FIGURE 3.**
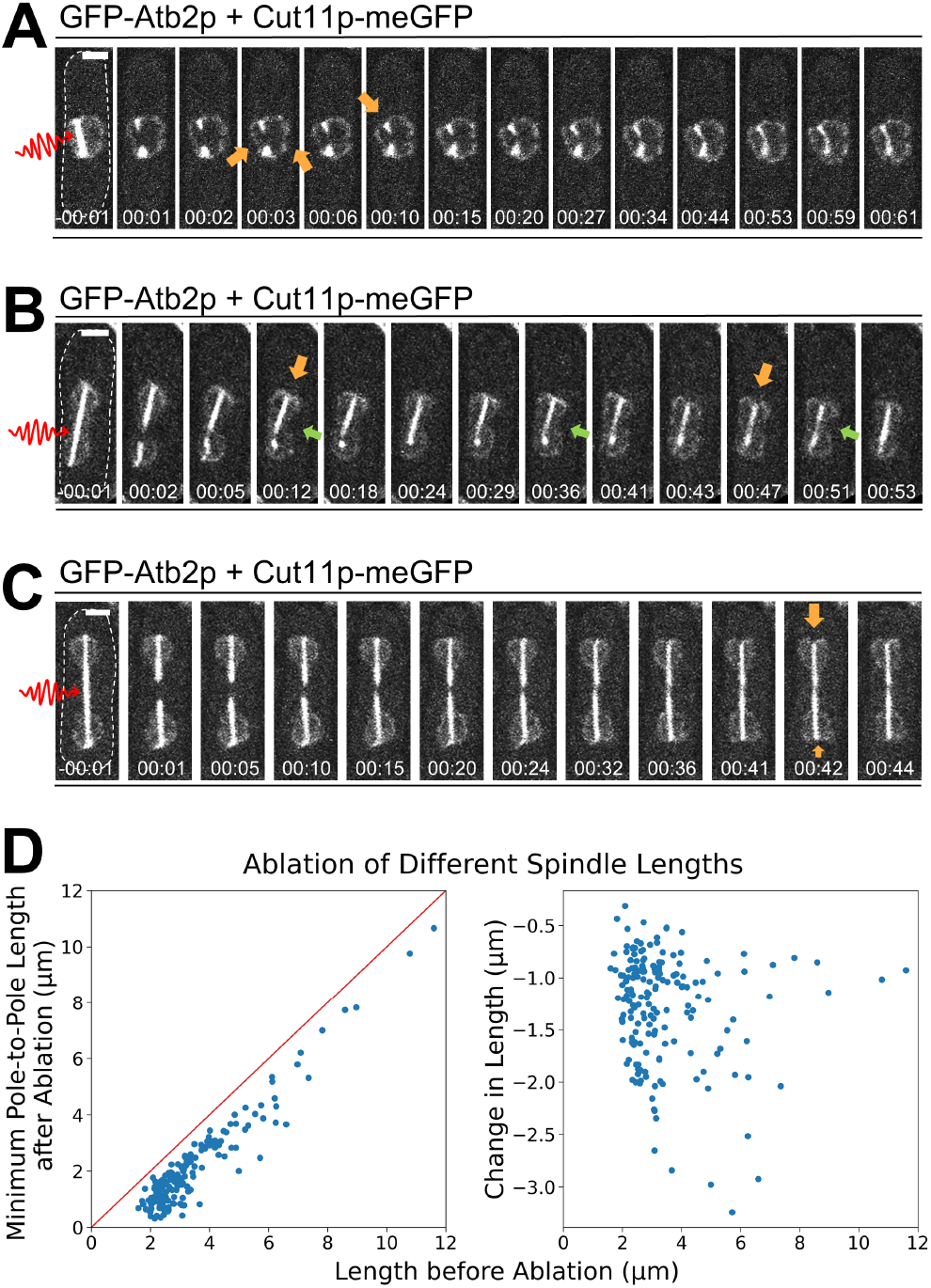
Pushing from the nuclear envelope on the spindle does not cause its collapse. (A)-(C) Three examples of *S. pombe* spindles expressing GFP-Atb2p and Cut11p-meGFP, ablated near the mid-region during phase two, mid-phase three, and late phase three of spindle elongation, respectively. Arrows mark the dents appearing at the nuclear envelope and cytoplasm boundary, indicating inward pulling forces after ablation. Scale bars, 2 *μ*m. Time, min:sec. Dashed line in first frame illustrates the cell boundary for clarity. In all three examples, inward dents of nuclear envelope appear at the ends of the spindles shortly after ablation (orange arrows). In (B), green arrows highlight that the nuclear envelope keeps its dumbbell shape after ablation. (D) Two scatter plots of *n* = 177 *S. pombe* control cells from Fig. 1 are shown. Left: scatter plot of minimum pole-to-pole length after ablation vs. length before ablation, along with a linear function showing y=x (red line). Right: scatter plot of collapsed length vs. length before ablation.

**FIGURE 4.**
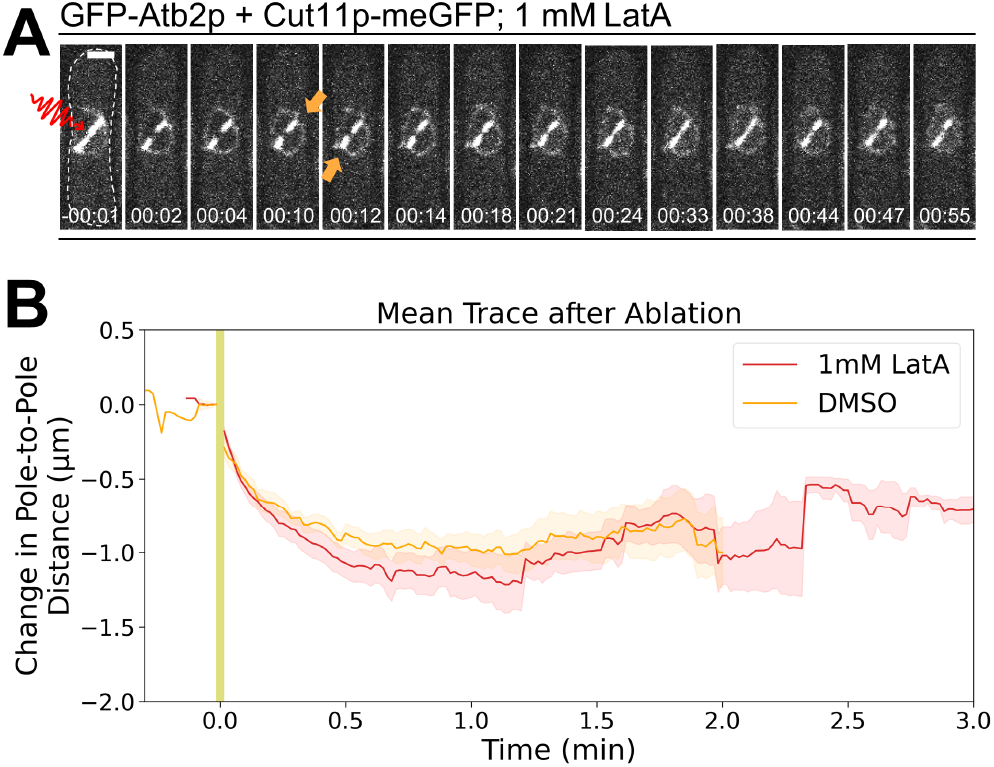
Actin depolymerization with latrunculin A does not affect spindle collapse. (A) An example of an ablated *S. pombe* spindle expressing GFP-Atb2p and Cut11p-meGFP treated with 1 mM latrunculin A during early phase three of spindle elongation. Arrows indicate nuclear envelope indentations following ablation. Dashed line indicates the cell membrane for clarity. Scale bar, 2 *μ*m. Time, min:sec. (B) The mean trace of change in pole-to-pole distance over time for *n* = 26 cells treated with 1 mM latrunculin A (red) is plotted along with a DMSO control (*n* = 9, orange). The shaded area shows the s.e.m. on each mean trace. We are unable to image during ablation, and the yellow shaded region indicates this period.

**FIGURE 5.**
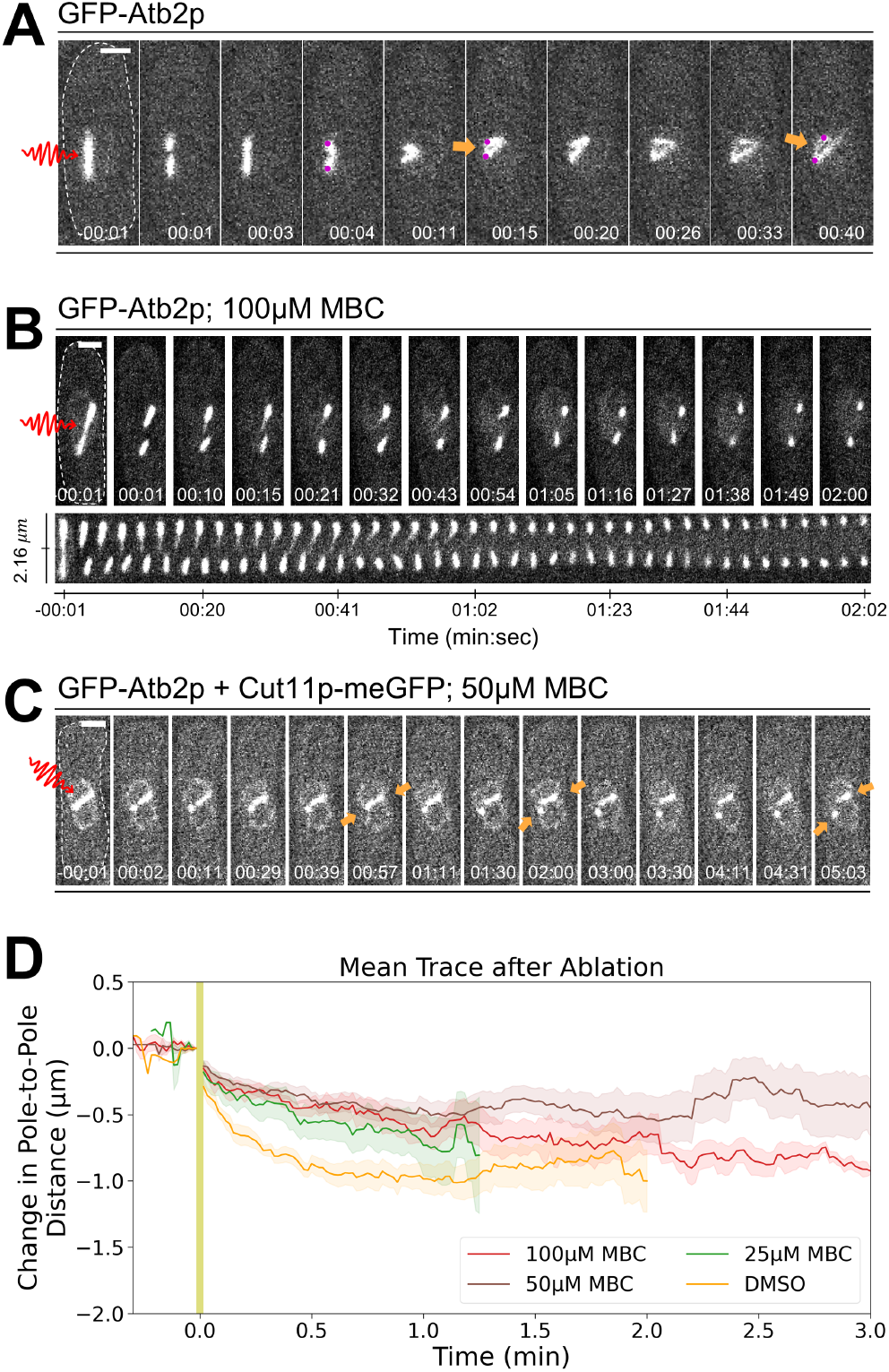
Spindle collapse after ablation requires microtubule dynamics. (A) Example *S. pombe* GFP-Atb2p spindle ablated near the mid-region during spindle elongation, showing connections of off-spindle-axis microtubules during its response. Arrows point to a bundle of microtubules. Circles (magenta) mark the supposed locations of spindle pole bodies. Example *S. pombe* spindle ablation in a cell treated with 100 *μ*M MBC, which does not lead to the spindle’s collapse. A montage of the spindle following ablation is displayed beneath. (C) An example of *S. pombe* spindle expressing GFP-Atb2p plus Cut11p-meGFP spindle and ablated near one pole in a cell treated with 50 *μ*M MBC during phase two of spindle elongation. The spindle collapses some but then seems to lose the connection between the poles. Arrows illustrate the changes in the nuclear envelope following ablation. Dashed line in the first frame illustrates the cell boundary for clarity. Scale bars, 2 *μ*m. Time, min:sec. (D) The mean trace of change in pole-to-pole distance over time for *n* = 7 cells treated with 100 *μ*M MBC (red), *n* = 23 cells treated with 50 *μ*M MBC (brown), and *n* = 8 cells treated with 25 *μ*M MBC (green), is shown along with a DMSO control (*n* = 9, orange). The shaded area shows the s.e.m. on each mean trace. We are unable to image during ablation, and the yellow shaded region indicates this period.

### Live cell spinning disk confocal fluorescent microscopy and laser ablation

We prepared a slide for imaging using a gelatin or agar pad on a microscope glass slide. We made gelatin pads by heating 125 mg of gelatin with 500 *μ*L of EMM5S at 90 °C for 5-10 minutes using a table-top dry heat bath. We made agar pads by melting YE + 1% agar in a microwave. In both cases, we pipetted a small amount onto a glass slide, topped it with a coverslip, and allowed it to cool. We, then, centrifuged 1 mL of culture at log phase (verified by measuring optical density) at 3000 RCF using a table-top centrifuge, decanted the supernatant, and resuspended the pellet in 20 *μ*L of media. We placed 5 *μ*L of resuspended culture on the agar or gelatin pad and covered it with a coverslip. Immediately before imaging, we sealed the coverslip using VALAP (1:1:1 Vase-line:lanolin:paraffin).

We performed live-cell spinning disc confocal imaging with laser ablation at room temperature of 22 °C as previously described [29, 30]. Briefly, we performed GFP imaging on a Nikon Ti-E stand on an Andor Dragonfly spinning disk confocal fluorescence microscope; spinning disk dichroic Chroma ZT405/488/561/640rpc; 488nm (50 mW) diode laser (75 ms to 120 ms exposures) with Borealis attachment (Andor); emission filter Chroma ET525/50m; and an Andor iXon3 camera. We performed imaging with a 100x 1.45 DIC Nikon objective and a 1.5x magnifier (built-in to the Dragonfly system). For imaging of mCherry, we used the same settings as above except for confocal excitation with 561 nm diode laser (Andor) and emission filter Chroma ET 500/50m. We collected frames every 1-3 s for the duration of imaging. We performed targeted laser ablation using an Andor Micropoint attachment to the above microscope with galvo-controlled steering to deliver 10-15 ns pulses at 20 Hz using a dye laser. We used two different laser dyes in these experiments with emission maxima at 625 nm in cells expressing GFP-Atb2p + Htap1-mCherry (MWE10) and 551 nm for all other strains (Andor). For software, we used Andor Fusion software to control acquisition and Andor IQ software to control the laser ablation system simultaneously.

### Analysis

We used ImageJ to crop and adjust brightness and contrast in all images. We also used ImageJ to convert the cropped.tif files to .avi for further analysis (see below). For brightness and contrast, we used linear adjustment and did not use interpolation or compression at any stage.

After the initial cropping and adjustment with ImageJ, we performed all further analyses using home-built python codes using the Jupyter notebook environment. Analysis code is available upon request. Our software loads in the cropped image stacks as .avi files. We then used our software to record the location of the distal (presumed spindle pole body) ends of the spindles by manual tracking. We calculated the end-to-end Cartesian distance over time between these two tracked ends. We averaged the end-to-end distance traces for cells of the same treatment condition to measure the dynamics of the collapse more precisely. To average traces acquired with 1 and 3 seconds intervals, we set the value of 3 seconds interval traces to be the same for 3 seconds. We calculated the mean trace by averaging points from each trace at that time point, and calculated the s.e.m. of all traces for each time point, and used it as an estimate on the error of this mean trace. We found the parameters, such as amplitude and *τ*, by fitting an exponential to the mean curve using the *curve_fit*() function in python.

The control mean trace of *n* = 177 cells in Fig. 1E comprises results from strains MWE2, MWE10, and MWE16 strains not treated with MBC or LatA. Stepwise dropoffs in Fig. 4 and Fig. 5 mean traces are caused by individual traces having different durations.

## RESULTS AND DISCUSSION

### *S. pombe* spindles collapse inward following laser ablation

To probe the mechanical stability and force balance of the mitotic spindle, we ablated spindles of live *S. pombe* cells expressing GFP-Atb2p (*α*-tubulin) (Fig. 1A). Consistent with previous reports [11, 12], we found that after severing the spindle in half via laser ablation, the fragments rapidly collapse toward each other (Fig. 1B). In most cases, the two spindle fragments collapsed directly toward each other along the spindle axis (Fig. 1B, Supp. Movie 1). We also occasionally observed the fragments to rotate within the nucleus as their poles moved toward each other (Fig. 1C).

While the collapse of *S. pombe* spindles in response to laser ablation has been observed [11, 12], previous work did not characterize the physical or molecular mechanism underlying it. To do so, we first quantified the collapse’s dynamics by tracking the end of each spindle fragment after ablation and calculating the distance between them. A typical example trace is shown in Fig. 1D, where we observe a sharp decrease in the pole-to-pole distance that begins immediately following ablation and continues for the next ~30 s. By averaging the trajectories for *n* = 177 ablated spindles, we determined that the average change in pole-to-pole distance over time follows an exponential relaxation response, as shown in Fig. 1E. This mean trace fits well to an exponential function with magnitude *A* = 0.84±0.02 *μ*m and time constant *τ* = 0.25±0.01 min (errors reported as standard deviations from the least-squares fit).

Previous work [11, 12] hypothesized that the spindle collapse in response to ablation might be caused by the viscoelastic relaxation of the nuclear envelope or chromosomes following disruption of extensile force from spindle elongation. Indeed, an exponential relaxation in response to ablation like the one we observe (Fig. 1E) is consistent with a passive viscoelastic response [31]. However, it is less clear that such a passive nuclear relaxation model could explain the spindle’s rotation within the nucleus, which we would not expect to be accompanied by an overall change in nuclear shape (Fig. 1C). Thus, we set out to test more directly whether forces from either rearrangement of the chromosomes within the nucleus or the nuclear envelope could explain spindle collapse.

### Passive viscoelastic relaxation of chromosomes does not cause the spindle’s collapse

Cohesin ensures sister chromatid cohesion and resists the poleward forces that pull the sister chromatids apart [32, 33]. When anaphase begins, cohesin is gradually degraded, allowing the sisters to separate and move to opposite spindle poles [34]. At the onset of stage three of spindle elongation, sufficient cohesin may remain to allow the chromosomes to oppose spindle elongation. Budding yeast chromosomal DNA has been shown to act as an entropic spring [35], and *S. pombe* chromosomes likely have similar material properties. We would thus expect a roughly spherical equilibrium conformation. During spindle elongation, the chromosomes are stretched out of this equilibrium state by the growing spindle. We considered whether spindle collapse might be driven by a mechanical relaxation of the chromosomes themselves to a more spherical confirmation.

To test this hypothesis, we ablated the spindle in cells expressing GFP-Atb2p and Hta1p-mCherry (histone H2A-*α*) to observe the simultaneous response of chromosomes and the spindle to ablation (Fig. 2). If the relaxation of chromosomal DNA powers spindle collapse, we would expect to observe chromosomal rearrangements accompanying the spindle’s movements.

However, following ablation, we could not detect any response of the histones in small spindles (before phase three of elongation). While bleaching of the Hta1p-mCherry signal by the ablation laser somewhat limits our observation immediately adjacent to the site of ablation, a detectable Hta1p-mCherry signal does remain following ablation (Fig. 2A and 2B). This signal did not significantly change shape in small spindles as the spindle collapsed. In other words, the spindle movements we observed seemed to occur within a largely stationary nucleus (Fig. 2A). The slight reorientation of the histones seen in Fig. 2A is not correlated with the inward movement of the collapsing spindle. While this observation is not wholly inconsistent with some chromosomal DNA movement, if significant rearrangement occurs, it must be very localized near the site of ablation (in the area that is bleached), and thus far from the translating spindle pole bodies.

In contrast, in longer spindles that had already entered phase three of spindle elongation before ablation, we did detect an inward histone movement that accompanied the collapsing spindle (Fig. 2B). Interestingly, as the spindle collapsed, inward indentations near the ends of the spindle fragments appeared in the histone localization (see arrows in Fig. 2B). This shape suggested inward pulling on the rest of the nucleus by the spindle during collapse rather than pushing on the spindle by the chromosomes. These spindle fragments then reconnected and resumed elongation.

Overall, these data do not support a strong role for viscoelastic relaxation of the chromosomes in underlying the collapse of either short or long ablated spindles. Therefore, we next tested whether the nuclear envelope might be pushing spindle poles together after ablation.

### Passive viscoelastic relaxation of the nuclear envelope does not cause the spindle’s collapse

During spindle elongation, the *S. pombe* nucleus grows and deforms to accommodate the lengthening spindle [20, 36, 37]. Pharmacological inhibition of fatty acid synthesis that prevents its growth can cause the nuclear envelope to exert sufficient force to bend or even break the spindle [20]. Thus, it is clear that the nuclear envelope is capable of exerting compressive stress on the spindle. To test whether relaxation of the nuclear envelope back toward a spherical shape might have a role in spindle collapse, we repeated the ablation experiment with a strain that expresses both GFP-Atb2p to mark the spindle and nuclear envelope marker Cut11p-meGFP [36] (Fig 3).

After ablation of small spindles, when the unperturbed nuclear envelope is not stretched and appears spherical, the collapsing spindle appeared to exert an inward pulling force on the nuclear envelope (Fig. 3A, Supp. Movie 2). This presumed inward force from the collapsing spindle induces a local inward curvature of the nuclear envelope. This shape change is not consistent with pushing from the nuclear envelope, so we conclude that the nuclear envelope does not power the collapse during stages one and two of mitosis, before the nucleus has begun to elongate.

We also considered whether the nuclear envelope might play a part in the collapse when the spindle has entered phase three of elongation and begun to increase the overall dimension of the nuclear envelope. If this were the case, we would expect the indentations shown in Fig. 3A to be absent in longer spindles, and that longer spindles would have a correspondingly greater magnitude of collapse due to the larger tension forces in the nuclear envelope at this stage.

In contrast to this prediction, indentations near the poles are still present in the nuclear envelope of spindles ablated late in phase three (Fig. 3B and C). Strikingly, in addition to indentations at the ends of spindle fragments, the nuclear envelope’s surface became less smooth and symmetric in some of these nuclei (Fig. 3B). Interestingly, in cells with a dumbbell-shaped nuclear envelope, the midsection’s curvature remained after ablation (green arrows, Fig. 3B). This behavior suggests mechanical isolation of this region from the rest of the nuclear envelope, rather than a uniform surface tension distributed around the entire envelope.

We also examined whether the extent of spindle collapse correlated with the pre-ablation spindle length. As illustrated in Fig. 3D, we found that the minimum pole-to-pole distance correlated with the pre-cut spindle length but that the extent of the collapse did not. That is, all spindles exhibit roughly the same degree of collapse, independent of their length.

In sum, we were unable to detect a role for nuclear envelope tension in spindle collapse at any stage of mitosis.

### Actin is not a factor in the collapse of the spindle

After finding that passive nuclear relaxation could not explain spindle collapse, we next investigated the role of the cytoskeleton. Depolymerizing actin delays anaphase onset by disrupting mitotic spindle stability [38], suggesting a possible role in spindle collapse. We examined whether actin was involved in spindle collapse by treating the cells expressing both GFP-Atb2p and Cut11p-meGFP with latrunculin A (LatA) (Fig. 4). We ablated cells within 2-4 minutes after the addition of LatA.

As shown in Fig. 4A, we observed a very similar collapse behavior in cells treated with 1 mM LatA as in untreated spindles. When we averaged the change in pole-to-pole distance over time for *n* = 26 spindles in cells treated with 1 mM LatA, we found that the magnitude and timescale of response were both very similar to control cells. These data suggest that actin is not a factor in spindle collapse.

### Microtubule dynamics are required for spindle collapse

We next tested whether the spindle itself might play an active force-generating role in spindle collapse. A rare observation shown in Fig. 5A and Supp. Movie 3 gave the first hint at a possible role for microtubules in the process. In this example, the spindle fragments first collapsed towards each other along the spindle’s main axis and then rotated. While rotating, the spindle fragments at the ablation site remained connected as each fragments’ end moved closer to the other. Shortly after, the GFP signal in the region between the ends increased (orange arrow, Fig. 5A), and a new bundle of microtubules began to appear between the fragments. This observation led us to investigate the role of microtubule dynamics in spindle collapse more closely.

To test whether spindle collapse required microtubule dynamics, we treated the cells with microtubule polymerization inhibitor methyl 2-benzimidazole carbamate (MBC) before ablation (Fig. 5B and 5C, Supp Movie 4). At 100 *μ*M MBC, few cells could maintain their spindles’ integrity, and all small spindles were depolymerized after the addition of MBC. However, we were still able to visualize longer spindles, which were more resistant to depolymerization. We ablated these spindles and examined their response to ablation (Fig. 5B). Spindle collapse was notably diminished in the presence of MBC. Instead of the fragments collapsing toward each other, they appeared to be free to undergo rotational diffusion.

We also examined the response to spindle ablation at 25 *μ*M and 50 *μ*M MBC, where we were still able to find stage two spindles. Interestingly, we sometimes observed spindles at these concentrations that seemed to initiate collapse, with the spindle fragments reconnecting as the spindle length initially shortened, but then appeared to become detached again and undergo diffusion apart from each other (Fig. 5C). Overall, however, collapse under this condition was lessened compared to untreated cells.

We quantified the pole-to-pole distance in ablated spindles at 25 *μ*M, 50 *μ*M, and 100 *μ*M MBC (Fig. 5D). In all cases, the mean spindle length does decrease over time, but it does so much more slowly and to a smaller extent than in control spindles, indicating that normal spindle collapse requires microtubule dynamics. The remaining collapse in the presence of MBC may still be microtubule-dependent, as we cannot completely inhibit microtubule dynamics without depolymerizing the spindle altogether.

### Minus-end directed motor dynein contributes to the collapse of the spindle

The microtubule dependence of spindle collapse led us to hypothesize a role for a minus-end directed microtubule motor, which might pull the spindle fragments together and shorten the pole-to-pole distance. *S. pombe*’s sole dynein heavy chain, dhc1, has known roles in meiosis but is not required for mitosis [25]. The evidence does suggest that it is involved in mitotic chromosome biorientation [26, 27], but with an unknown mechanism. It was also an interesting candidate for its known roles in maintaining spindle bipolarity and local force balance in higher eukaryotes [23, 24, 39, 40].

We examined the collapse response in *S. pombe* cells with dhc1 truncated after residue 1890 [41], which we refer to as dhc1ΔC. Spindle collapse was strikingly disrupted in dhc1ΔC cells (Fig. 6, Supp. Movie 5). As with MBC treatment, the fragments of ablated spindles in dhc1ΔC cells often showed a prolonged diffusive search rather than a collapse (Fig. 6A and B). We also noted the lack of any further collapse at the onset of a new connection compared to wild-type spindles. On quantification, we found that the pole-to-pole distance in dhc1ΔC shows a much smaller decrease following ablation than control spindles (Fig. 6C). Thus, dynein plays an important role in powering the collapse of the *S. pombe* spindle in response to ablation. We speculate that this role may be through minus-end directed dynein motor activity that slides antiparallel microtubules to bring SPBs closer together following ablation.

**FIGURE 6.**
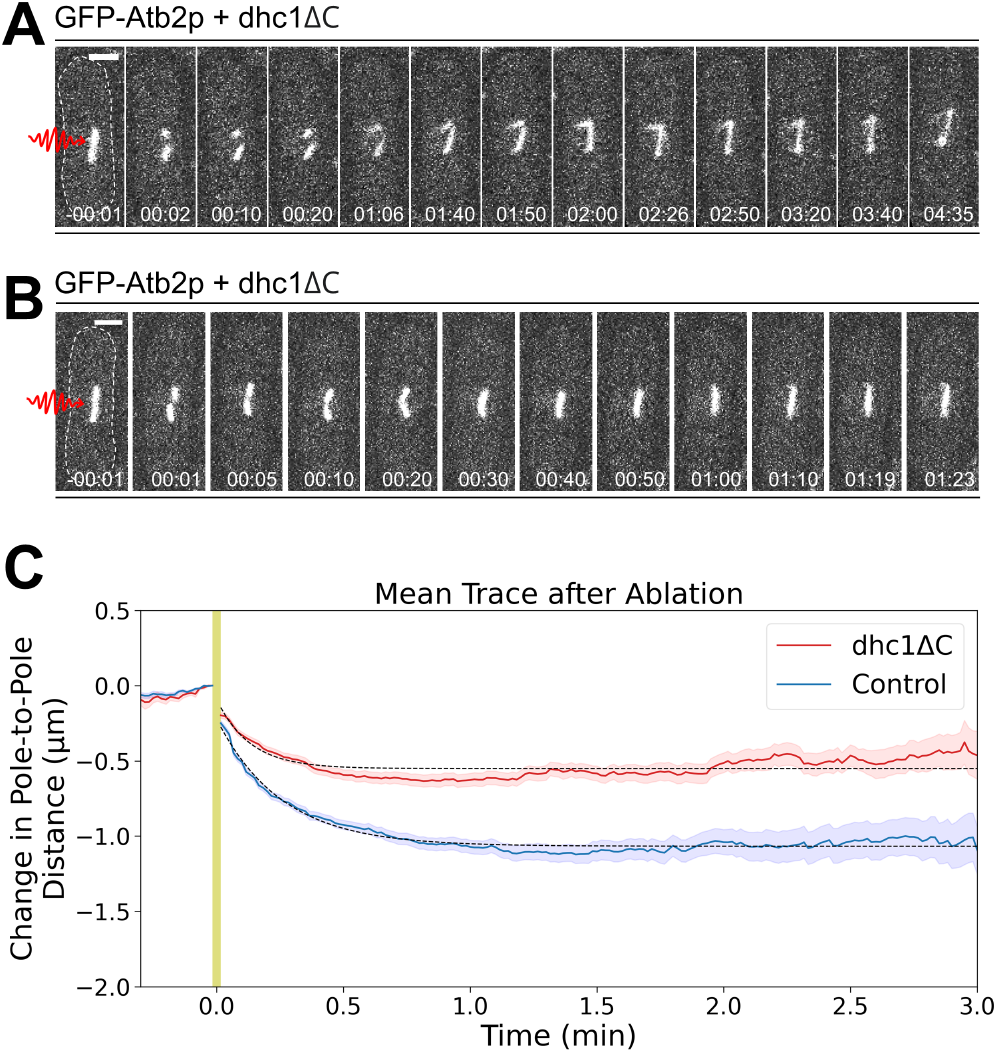
Spindles in dhc1ΔC cells exhibit reduced collapse following ablation. (A) and (B) Two example *S. pombe* GFP-Atb2p and dhc1ΔC spindles ablated near the mid-region during phase two of spindle elongation. Dashed line in the first frame illustrates the cell boundary for clarity. Scale bars, 2 *μ*m. Time, min:sec. (C) The mean trace of change in pole-to-pole distance over time for *n* = 72 ablated spindles in GFP-Atb2p plus dhc1ΔC cells is plotted (solid red line). Black dotted line is a fit to an exponential, with amplitude 0.05 *μ*m and time constant 0.13 0.02 minute. We also plot the analogous mean trace and exponential fit for *n* = 177 control untreated cells from Fig. 1 for comparison (blue). We are unable to image during ablation, and the yellow shaded region indicates this period.

## CONCLUSIONS

Here, we have characterized the physical mechanism of the collapse of the *S. pombe* spindle in response to laser ablation. The two fragments of severed spindles collapse toward each other, and their trajectory follows an exponential curve. The character of this response initially seemed consistent with viscoelastic relaxation, as was previously suggested. However, we find that neither the chromosomes nor the nuclear envelope respond in a way to suggest that they power the spindle collapse. Actin depolymerization also does not affect the collapse. Instead, we find that microtubule polymerization is required for collapse. C-terminal truncation of the minus-end directed microtubule motor dynein also disrupts spindle collapse.

While the role of dynein in spindle collapse is clear, we do note that dhc1ΔC cells still demonstrate some degree of spindle collapse, so it must not be the only contributor. There are a few potential explanations for this result. First, it may be that there is redundancy between multiple minus-end directed microtubule motors, of which dynein is not the only example in *S. pombe*. Alternatively, we cannot fully exclude the possibility that viscoelastic relaxation of the nucleus or nuclear envelope, while not primarily responsible for it, may make some contribution to the collapse.

In future work, it will be interesting to further examine the potential of involvement of additional minus-end directed microtubule motors, or of other microtubule crosslinkers. In addition, the behavior of the nuclear envelope in response to ablation raises important questions about the mechanics of the nuclear envelope and how it is shaped by its physical connection to the spindle.

## AUTHOR CONTRIBUTIONS

PZ and MWE designed experiments. PZ performed experiments and wrote analysis software. PZ and MWE performed analysis and wrote the manuscript.

## ACKNOWLEDGEMENTS

We thank Eva Johannes and Mariusz Zareba of the NC State Cellular and Molecular Imaging Facility, Kerry Bloom, Arthur Molines, Kimberly Bellingham-Johnstun and members of the Elting lab for helpful discussions, and we thank Caroline Laplante and Fred Chang for helpful discussions and for providing strains. PZ acknowledges support by the Provost’s Professional Experience Program, the Office of Undergraduate Research at North Carolina State University, and the National Society of Physics Students. MWE acknowledges support by NIH 1R35GM138083 and NSF 1935260.

## SUPPORTING MATERIAL

### Supplemental Movie Legends

**Supplemental Movie 1**: Timelapse movie of Fig. 1B. An example *S. pombe* GFP-Atb2p spindle ablated near the mid-region during phase two of spindle elongation. Spindle ends (yellow arrows) collapse toward each other after ablation. Star (magenta) marks the ablation point. Time, min:sec. Images were taken at 3 sec intervals. Note that there is a gap in frames as we are unable to image during ablation.

**Supplemental Movie 2**: Timelapse movie of Fig. 3A. An example S. pombe spindle expressing GFP-Atb2p and Cut11p-meGFP (which labels the nuclear envelope protein), ablated near the mid-region during phase two of spindle elongation. Arrows (yellow) mark the dents appearing at the nuclear envelope and cytoplasm boundary, indicating inward pulling forces after ablation. Star (magenta) marks the ablation point. Time, min:sec. Images were taken at 1 sec intervals. Note that there is a gap in frames as we are unable to image during ablation.

**Supplemental Movie 3**: Timelapse movie of Fig. 5A. An example *S. pombe* GFP-Atb2p spindle ablated near the mid-region during phase two of spindle elongation. Arrows (yellow) point to a bundle of microtubules. Star (magenta) marks the ablation point. Time, min:sec. Images were taken at 1 sec intervals. Note that there is a gap in frames as we are unable to image during ablation.

**Supplemental Movie 4**: Timelapse movie of Fig. 5B. An example *S. pombe* GFP-Atb2p spindle ablated near the mid-region during mid-phase three of spindle elongation. Star (magenta) marks the ablation point. Spindle ablation in a cell treated with 100*μ*M MBC does not lead to the collapse of the spindle. Time, min:sec. Images were taken at 1 sec intervals. Note that there is a gap in frames as we are unable to image during ablation.

**Supplemental Movie 5**: Timelapse movie of Fig. 6A. An example of *S. pombe* GFP-Atb2p plus dhc1ΔC spindle ablated near the mid-region during phase two of spindle elongation. Star (magenta) marks the ablation point. Time, min:sec. Images were taken at 1 sec intervals. Note that there is a gap in frames as we are unable to image during ablation.

## Supplemental Table

**TABLE S1.** Fission yeast strains used in this study

| Strains | Genotypes | Figures used | Sources |
| --- | --- | --- | --- |
| MWE2 | *h+ GFP-atb2:kanMX ade6- leu1-32 ura4-D18* | 1B-C,5A-B | FC2861, Chang lab stock |
| MWE10 | *h+ hta1-mCherry:kanMX GFP-atb2:kanMX ade6- leu1-32 ura4-* | 2A-B | FC2859, Chang lab stock. Original source HY1212. |
| MWE16 | *h+ nmt41-GFP-atb2:hpnMX6 + cut11-meGFP:kanMX6* | 3A-C,4A,5C | CL579-1, gift of Caroline Laplante |
| MWE23 | *dhc1-aa1890-2527::kanMX leu1-32::SV40:GFP-atb2:leu1+ ura4- ade6-* | 6A-B | FC2021, Chang lab stock. Original source SB636 [41]) |

## Notes

### Competing Interest Statement

The authors have declared no competing interest.

## REFERENCES

[1] Santaguida, S., and A. Amon, 2015. Short- and long-term effects of chromosome mis-segregation and aneuploidy. Nature Reviews. Molecular Cell Biology 16:473–485.

[2] Singh, V. P., and J. L. Gerton, 2015. Cohesin and human disease: lessons from mouse models. Current Opinion in Cell Biology 37:9–9. http://www.sciencedirect.com/science/article/pii/S0955067415001088.

[3] Sawin, K. E., and P. T. Tran, 2006. Cytoplasmic microtubule organization in fission yeast. Yeast 23:1001–1014.

[4] Tolić-Nørrelykke, I. M., 2010. Force and length regulation in the microtubule cytoskeleton: lessons from fission yeast. Curr. Opin. Cell Biol. 22:21–28.

[5] Ding, R., K. L. McDonald, and J. R. McIntosh, 1993. Three-dimensional reconstruction and analysis of mitotic spindles from the yeast, Schizosaccharomyces pombe. Journal of Cell Biology 120:141–141. https://rupress.org/jcb/article/120/1/141/14563/Three-dimensional-reconstruction-and-analysis-of, publisher: The Rockefeller University Press.

[6] Mitchison, J. M., 1970. Chapter 7 Physiological and Cytological Methods for Schizosaccharomyces pombe. In D. M. Prescott, editor, Methods in Cell Biology, Academic Press, volume 4, 131–165.

[7] Hagan, I. M., and J. S. Hyams, 1988. The use of cell division cycle mutants to investigate the control of microtubule distribution in the fission yeast Schizosaccharomyces pombe. J. Cell Sci. 89 (Pt 3):343–357.

[8] Nabeshima, K., T. Nakagawa, A. F. Straight, A. Murray, Y. Chikashige, Y. M. Yamashita, Y. Hiraoka, and M. Yanagida, 1998. Dynamics of Centromeres during Metaphase–Anaphase Transition in Fission Yeast: Dis1 Is Implicated in Force Balance in Metaphase Bipolar Spindle. Molecular Biology of the Cell 9:3211–3225. https://www.molbiolcell.org/doi/10.1091/mbc.9.11.3211.

[9] Sagolla, M. J., S. Uzawa, and W. Z. Cande, 2003. Individual microtubule dynamics contribute to the function of mitotic and cytoplasmic arrays in fission yeast. J. Cell Sci. 116:4891–4903.

[10] Oliferenko, S., and M. K. Balasubramanian, 2002. Astral microtubules monitor metaphase spindle alignment in fission yeast. Nat. Cell Biol. 4:816–820.

[11] Tolić-Nørrelykke, I. M., L. Sacconi, G. Thon, and F. S. Pavone, 2004. Positioning and Elongation of the Fission Yeast Spindle by Microtubule-Based Pushing. Current Biology 14:1181–1181. http://www.sciencedirect.com/science/article/pii/S0960982204004257.

[12] Khodjakov, A., S. La Terra, and F. Chang, 2004. Laser Microsurgery in Fission Yeast: Role of the Mitotic Spindle Midzone in Anaphase B. Current Biology 14:1330–1330. http://www.sciencedirect.com/science/article/pii/S096098220400510X.

[13] McIntosh, J. R., and E. T. O’Toole, 1999. Life cycles of yeast spindle pole bodies: Getting microtubules into a closed nucleus. Biology of the Cell 91:305–312. http://doi.wiley.com/10.1111/j.1768-322X.1999.tb01089.x.

[14] Tanaka, K., and T. Kanbe, 1986. Mitosis in the fission yeast Schizosaccharomyces pombe as revealed by freeze-substitution electron microscopy. J. Cell Sci. 80:253–268.

[15] Masuda, H., T. Hirano, M. Yanagida, and W. Z. Cande, 1990. In vitro reactivation of spindle elongation in fission yeast nuc2 mutant cells. The Journal of Cell Biology 110:417–417. https://rupress.org/jcb/article/110/2/417/55727/In-vitro-reactivation-of-spindle-elongation-in.

[16] Mallavarapu, A., K. Sawin, and T. Mitchison, 1999. A switch in microtubule dynamics at the onset of anaphase B in the mitotic spindle of Schizosaccharomyces pombe. Current Biology 9:1423–1423. http://www.sciencedirect.com/science/article/pii/S0960982200800901.

[17] Hagan, I., and M. Yanagida, 1992. Kinesin-related cut7 protein associates with mitotic and meiotic spindles in fission yeast. Nature 356:74–76.

[18] Fu, C., J. J. Ward, I. Loiodice, G. Velve-Casquillas, F. J. Nedelec, and P. T. Tran, 2009. Phospho-Regulated Interaction between Kinesin-6 Klp9p and Microtubule Bundler Ase1p Promotes Spindle Elongation. Dev. Cell 17:257–267.

[19] Simeonov, D. R., K. Kenny, L. Seo, A. Moyer, J. Allen, and J. L. Paluh, 2009. Distinct Kinesin-14 mitotic mechanisms in spindle bipolarity. Cell Cycle 8:3571–3583.

[20] Yam, C., Y. He, D. Zhang, K.-H. Chiam, and S. Oliferenko, 2011. Divergent Strategies for Controlling the Nuclear Membrane Satisfy Geometric Constraints during Nuclear Division. Current Biology 21:1314–1314. http://www.sciencedirect.com/science/article/pii/S0960982211007251.

[21] Pavin, N., and I. M. Tolić, 2016. Self-Organization and Forces in the Mitotic Spindle. Annu. Rev. Biophys. 45:279–298.

[22] Dumont, S., and T. J. Mitchison, 2009. Force and length in the mitotic spindle. Curr. Biol. 19:R749–61.

[23] Verde, F., J. M. Berrez, C. Antony, and E. Karsenti, 1991. Taxol-induced microtubule asters in mitotic extracts of Xenopus eggs: requirement for phosphorylated factors and cytoplasmic dynein. J. Cell Biol. 112:1177–1187.

[24] Gordon, M. B., L. Howard, and D. A. Compton, 2001. Chromosome movement in mitosis requires microtubule anchorage at spindle poles. J. Cell Biol. 152:425–434.

[25] Yamamoto, A., R. R. West, J. R. McIntosh, and Y. Hiraoka, 1999. A Cytoplasmic Dynein Heavy Chain Is Required for Oscillatory Nuclear Movement of Meiotic Prophase and Efficient Meiotic Recombination in Fission Yeast. The Journal of Cell Biology 145:1233–1250. https://www.ncbi.nlm.nih.gov/pmc/articles/PMC2133150/.

[26] Grishchuk, E. L., I. S. Spiridonov, and J. R. McIntosh, 2007. Mitotic chromosome biorientation in fission yeast is enhanced by dynein and a minus-end-directed, kinesin-like protein. Molecular Biology of the Cell 18:2216–2225.

[27] Courtheoux, T., G. Gay, C. Reyes, S. Goldstone, Y. Gachet, and S. Tournier, 2007. Dynein participates in chromosome segregation in fission yeast. Biology of the Cell 99:627–637.

[28] Forsburg, S. L., and N. Rhind, 2006. Basic Methods for Fission Yeast. Yeast (Chichester, England) 23:173–173. https://www.ncbi.nlm.nih.gov/pmc/articles/PMC5074380/.

[29] Elting, M. W., M. Prakash, D. B. Udy, and S. Dumont, 2017. Mapping Load-Bearing in the Mammalian Spindle Reveals Local Kinetochore Fiber Anchorage that Provides Mechanical Isolation and Redundancy. Current Biology 27:2112–2122.e5. http://www.sciencedirect.com/science/article/pii/S0960982217307200.

[30] Begley, M. A., A. L. Solon, E. M. Davis, M. G. Sherrill, R. Ohi, and M. W. Elting, 2020. K-fiber bundles in the mitotic spindle are mechanically reinforced by Kif15. bioRxiv 2020.05.19.104661. https://www.biorxiv.org/content/10.1101/2020.05.19.104661v1.

[31] Roca-Cusachs, P., V. Conte, and X. Trepat, 2017. Quantifying forces in cell biology. Nat. Cell Biol. 19:742–751.

[32] Bernard, P., J. F. Maure, J. F. Partridge, S. Genier, J. P. Javerzat, and R. C. Allshire, 2001. Requirement of heterochromatin for cohesion at centromeres. Science 294:2539–2542.

[33] Gerton, J. L., 2007. Enhancing togetherness: kinetochores and cohesion. Genes & Development 21:238–238. http://www.genesdev.org/cgi/doi/10.1101/gad.1523107.

[34] Tomonaga, T., K. Nagao, Y. Kawasaki, K. Furuya, A. Murakami, J. Morishita, T. Yuasa, T. Sutani, S. E. Kearsey, F. Uhlmann, K. Nasmyth, and M. Yanagida, 2000. Characterization of fission yeast cohesin: essential anaphase proteolysis of Rad21 phosphorylated in the S phase. Genes Dev. 14:2757–2770.

[35] Vasquez, P. A., C. Hult, D. Adalsteinsson, J. Lawrimore, M. G. Forest, and K. Bloom, 2016. Entropy gives rise to topologically associating domains. Nucleic Acids Research 44:5540–5549.

[36] West, R. R., E. V. Vaisberg, R. Ding, P. Nurse, and J. R. McIntosh, 1998. Cut11 +: A gene required for cell cycle-dependent spindle pole body anchoring in the nuclear envelope and bipolar spindle formation in Schizosaccharomyces pombe. Mol. Biol. Cell 9:2839–2855.

[37] Saitoh, S., K. Takahashi, K. Nabeshima, Y. Yamashita, Y. Nakaseko, A. Hirata, and M. Yanagida, 1996. Aberrant mitosis in fission yeast mutants defective in fatty acid synthetase and acetyl CoA carboxylase. J. Cell Biol. 134:949–961.

[38] Meadows, J. C., and J. Millar, 2008. Latrunculin A Delays Anaphase Onset in Fission Yeast by Disrupting an Ase1-independent Pathway Controlling Mitotic Spindle Stability. Molecular Biology of the Cell 19:3713–3713. https://www.molbiolcell.org/doi/10.1091/mbc.e08-02-0164.

[39] Elting, M. W., C. L. Hueschen, D. B. Udy, and S. Dumont, 2014. Force on spindle microtubule minus ends moves chromosomes. Journal of Cell Biology 206:245–245. https://rupress.org/jcb/article/206/2/245/37798/Force-on-spindle-microtubule-minus-ends-moves, publisher: The Rockefeller University Press.

[40] Sikirzhytski, V., V. Magidson, J. B. Steinman, J. He, M. Le Berre, I. Tikhonenko, J. G. Ault, B. F. McEwen, J. K. Chen, H. Sui, M. Piel, T. M. Kapoor, and A. Khodjakov, 2014. Direct kinetochore-spindle pole connections are not required for chromosome segregation. J. Cell Biol. 206:231–243.

[41] Bratman, S. V., and F. Chang, 2007. Stabilization of overlapping microtubules by fission yeast CLASP. Developmental Cell 13:812–827.

